# AstraBIND: Graph Attention Network for Predicting Ligand Binding Sites

**DOI:** 10.1101/2025.11.10.687555

**Authors:** Aniruddh Goteti, Alexandra Vasilyeva, Çağlar Bozkurt

## Abstract

Predicting ligand binding sites is central to computational biology and drug discovery. Existing machine learning approaches either use protein sequence, structure, or both. While structure-based deep learning models typically outperform sequence-based methods, they often require high computational cost or ligand-specific data, forcing a trade-off between accuracy and scalability.

We present AstraBIND, a lightweight graph neural network that bridges this gap by integrating protein sequence, structure (experimental or predicted), and homology information to predict ligand classes and binding residues within minutes. The model employs a GATv2 architecture with 0.9 M parameters, trained on ¿250 000 curated protein–ligand complexes across 16 ligand categories. By encoding residue-level features and spatial geometry through graph attention, AstraBIND identifies binding residues and ligand types while maintaining structural consistency.

In benchmarking, AstraBIND achieved a weighted macro-F1 of 0.47 across all ligand classes, with top performance for nucleotides (F1 = 0.79), porphyrins (0.74), and cofactors (0.73). Case studies, including p53 and CRFR1, demonstrate robust pocket localization for diverse proteins. Combined with its minimal inference time and broad ligand coverage, AstraBIND enables rapid in-silico screening and integration into laboratory workflows. Together with other Astra ML models (1; 2), it represents a step toward real-time protein design and validation pipelines.

Astra models are available at https://www.orbion.life.

## Introduction

Prediction of ligand binding sites for a protein is a crucial task in computational biology, in particular in the area of drug development. Over the past years, multiple machine learning (ML) methods have been developed to predict binding sites based on the protein’s sequence (BIRDS, Pseq2Sites, Seq-InSite), structure (DeepPocket, PURes-Net, PGpocket, GrASP, VN-EGNN) or both (DELIA, T5-GAT Ensemble, IF-SitePred). Overall, structure-based deep learning approaches, such as 3D convolutional neural networks (CNNs), U-Net segmentation models and equivariant graph neural networks, have surpassed classical geometry-based tools in reliably identifying pockets and active sites. Precision can be improved further, but only for known specific ligands or at a high computational cost (e.g. leveraging molecular dynamics simulations). At the same time, sequence-based models offer orders-of-magnitude faster predictions, but at a trade-off with accuracy.

To address this gap, we have developed AstraBIND - an ML model that accepts protein sequence as input, enhances it with an experimental or predicted structure and a homology search, and outputs predicted ligands and their binding sites in just a few minutes per protein. AstraBIND is a Graph Attention Network (GATv2) with 892,445 trainable parameters; by training on curated ligand binding databases and leveraging the graph architecture to process protein information on a structure voxel level, AstraBIND can screen the input protein against a broad library of ligands and identify the residues that are likely involved in ligand binding, as well as the ligand type. Model output consists of predicted ligand IDs with corresponding protein residue numbers, with the graph architecture ensuring that the spatial logic is maintained, i.e. only residues that are predicted to be close together participate in the same interaction. Importantly, AstraBIND offers state-of-the-art accuracy combined with fast processing, which makes it suitable for integration within laboratory pipelines along with other Astra ML models.(1; 2)

## Methods

### Dataset structure

For Astra model training, experimentally supported protein-ligand binding datasets from publicly available curated databases were selected, leading to a dataset of *>*250k protein-ligand interactions. Receptor and ligand structures were saved as local PDB files (managed by a local index for fast lookup). Binding residues were then parsed into integer residue numbers that preserve original PDB indexing. Ligand identifiers were mapped to 16 consistent categories to provide graph level labels that match how the model reasons about ligands:

- Peptide
- Metal ions
- Metal complexes
- Nucleotides & analogs
- Nucleosides
- Cofactors
- Carbohydrates
- Amino acids
- Vitamins
- Lipids
- Organic acids
- Steroids
- Pharmaceuticals
- Solvents & small molecules
- Porphyrins
- Other

The processed training set yielded the protein graphs with residue level binding labels, residue level pocket IDs and a graph level ligand class per complex. Node representation (per residue) includes:

- **Amino acid identity** as a 21 channel one hot encoding (standard 20 plus unknown).
- **Physicochemical properties** (13 channels) capturing hydrophobicity, polarity, charge, pKa like signals, and simple size proxies.
- **Structural values** from the CA atom: normalized x, y, z coordinates and B factor.
- **Ligand interaction features** (5 channels) computed when ligand atoms are available: distance to ligand centroid, minimum distance to any ligand atom, a contact indicator within 6 Å, an inverse distance interaction strength, and two coarse interaction potentials (hydrogen bond potential within 3.5 Å for polar residues; hydrophobic contact potential within 4.5 Å for hydrophobic residues).

Residue–residue edges are created when CA–CA distance is within 8 Å (undirected connectivity implemented as two directed edges). Each directed edge carries 7 attributes: Euclidean distance, inverse distance, sequence separation, a sequential neighbour flag, normalized ligand distance for each endpoint residue, and a joint near ligand flag indicating both residues are close to the ligand. Graph metadata includes original PDB residue numbering, a ligand category string, simple statistics (binding ratio, protein size) and a PDB like identifier. Binding residues are clustered into pockets; each residue receives a pocket ID, capped at 8 unique pockets to maintain a stable classification target.

### Model architecture: multi-task GNN with ligand awareness

A stack of graph attention layers (GATv2) with edge attributes encodes the residue graph. Attention over edges lets the model weigh structural neighbours while using the 7 dimensional edge features to modulate message passing by geometry and ligand proximity. Three types of heads were used:

- **Binding site head (node level)**: a binary classifier predicts binding vs non binding per residue.
- **Pocket head (node level)**: a multiclass classifier assigns pocket IDs (up to 8).
- **Ligand heads (graph level)**: a two stage hierarchy improves ligand recognition a) A coarse three way ligand classifier provides a stabilizing prior. b) A fine grained ligand classifier predicts a richer ligand class set. The coarse output is mapped through a learned projection and added as a bias to the fine logits. c) The pooled graph embedding is softly gated by the average binding probability across residues, so ligand predictions activate mainly when binding evidence exists at the residue level.

Binding is localized at residues, while ligand identity is a global property of the protein–ligand situation. Gating the ligand head by binding signals reduces spurious ligand predictions on structures with no credible binding evidence. The hierarchical coarse→fine mapping improves rare class stability and generalization.

### Training workflow

#### Input assembly

1. Read data training TSV (includes ligand info and binding residue strings).
2. Locate receptor and ligand PDB files in local directories via an indexer for fast, robust file access.
3. Parse binding residues into integer indices (matching original PDB numbering).
4. When ligand atoms are available, compute ligand interaction features per residue.

#### Graph building

1. Create a single residue graph per receptor–ligand complex with node, edge, and metadata features as described above.
2. Add labels: residue binding flag, residue pocket ID, and graph level ligand category.

#### Optimization

1. Train all three tasks jointly (binding, pockets, ligand class).
2. Handle imbalance with architecture choices (binding gated ligand head, capped pocket classes) and with evaluation thresholds tuned to binding detection quality.
3. Save the final checkpoint.

### Inference

#### Sequence to structure

1. Perform a homology search to identify PDB structures and UniProt matches.
2. Choose a primary structure from top PDB hits; if unavailable, fall back to the AlphaFold model. This guarantees a structure for feature extraction even without an experimental complex.

#### Contextual ligand discovery

1. Experimental ligands: extract ligands present in the chosen structure and build context aware graphs for each ligand.
2. Panel ligands: assemble a broader ligand panel from external data sources aligned to the best UniProt ID and homology (pre integrated in the app). Each candidate carries type and evidence metadata.
3. General context: build a ligand agnostic graph that lets the model detect binding regions purely from protein geometry.

#### Graph creation per context

1. Parse receptor residues from the structure and compute residue features.
2. If ligand coordinates or a ligand structure are available, compute ligand interaction features to enrich residues near the ligand.
3. Apply the same ligand category mapping used during training to keep category semantics aligned.

#### Model forward pass

1. Batch the graph and run the GNN to obtain residue binding logits, residue pocket logits, and graph level ligand logits.
2. Convert residue binding probabilities into compact binding regions via clustering, optionally gated with a ligand distance window (for example, using 12 Å as a gating radius when ligand coordinates are known).

**Merging and ranking**

1. Merge regions from all contexts (experimental, panel, general).
2. Score regions by combining strength of binding signal, pocket coherence, and ligand plausibility.
3. Return the top regions with residue ranges in original numbering, confidence scores, qualitative labels, and evidence tags (experimental structure vs panel vs general model pass).
4. Optionally filter by a minimum confidence threshold before returning to the client.

#### Operational interface

1. Asynchronous endpoint submits jobs to a queue and returns a job ID for progress polling.
2. A synchronous endpoint exists for convenience; it internally uses the same queue but waits for completion.
3. Health and utility endpoints report model loading status and available ligand categories.

### User Experience

#### End-to-end flow

AstraBIND is integrated into the Orbion web platform. Users provide a protein sequence or UniProt ID (optionally selecting ligand categories of interest). The app performs a homology search and retrieves a structure (PDB if available, otherwise AlphaFold2), builds graph contexts (experimental ligands, panel ligands, and ligand-agnostic geometry), and runs the GATv2 encoder to produce *residue-level binding probabilities, pocket assignments*, and *graph-level ligand class scores*. Outputs are returned as ranked binding regions with confidence values and evidence tags (experimental / panel / general), and rendered in synchronized 2D and 3D views. Thresholds can be viewed in *calibrated* and *fixed* modes.

#### 2D & 3D visualization

The 2D view displays a per-residue track with binding probability (heatmap) and pocket IDs (color bands), plus chips for top ligand class predictions per region. Users can filter by ligand category, toggle confidence thresholds, and inspect evidence provenance. The 3D viewer (implemented using the `3Dmol.js`library)(17) overlays predicted binding residues and pocket surfaces onto the structure; when ligand coordinates are present, candidate ligands are shown as sticks/spheres. Tooltips expose `(residue number, residue name, pocket ID, binding probability, ligand class, evidence)`. The linked views allow quick scanning along sequence and immediate spatial reasoning in the folded structure.

## Results

### Performance Metrics

The model showed positive results on hold-out dataset testing, with weighted macro-F1 of 0.47 (see Table 1). Among the 16 ligand categories, highest F1 was seen for nucleotides (F1=0.79), porphyrins (F1=0.74) and cofactors (F1=0.73). On the other hand, several labels (lipid, nucleoside, organic acid, other, peptide, steroid and vitamin) showed low performance (F1=0.1 or less). For all of these labels except “Other”, support was low (from 5 to 224), indicating that poor performance was driven by low sample numbers in these categories.

**Table 1.** Key model performance results on hold-out testing for 16 ligand categories.

| Label | Precision | Recall | F1 | Support |
| --- | --- | --- | --- | --- |
| Amino acid | 0.2 | 0.35 | 0.25 | 498 |
| Carbohydrate | 0.44 | 0.26 | 0.32 | 1659 |
| Cofactor | 0.73 | 0.73 | 0.73 | 2396 |
| Lipid | 0.06 | 0.6 | 0.1 | 199 |
| Metal complex | 0.24 | 0.86 | 0.38 | 1553 |
| Metal ion | 0.98 | 0.46 | 0.62 | 10200 |
| Nucleoside | 0.02 | 0.33 | 0.03 | 87 |
| Nucleotide | 0.94 | 0.68 | 0.79 | 6424 |
| Organic acid | 0.04 | 0.21 | 0.07 | 224 |
| Other | 0.76 | 0 | 0.01 | 8498 |
| Peptide | 0.06 | 0.63 | 0.1 | 137 |
| Pharmaceutical | 0.44 | 0.69 | 0.54 | 1742 |
| Porphyrin | 0.62 | 0.91 | 0.74 | 1452 |
| Solvent | 0.26 | 0.58 | 0.36 | 1280 |
| Steroid | 0 | 0.2 | 0 | 5 |
| Vitamin | 0.06 | 0.53 | 0.1 | 102 |
| Macro avg | 0.36 | 0.5 | 0.32 | 36456 |
| Weighted avg | 0.75 | 0.45 | 0.47 | 36456 |

### Case Studies

Performance of the AstraBIND model in identifying binding sites can best be showcased with example case studies (see Figure 2). Overall, model performance was satisfactory for relatively small and soluble proteins: the key DNA-binding chain of p53 was successfully identified, and residues close to relevant binding pockets were highlighted for lysozyme C. However, for some of the more complicated proteins binding pocket location was not identified: for example, for corticotropin-releasing factor receptor 1 (CRFR1) a known small molecule ligand was correctly identified (0JS, or 8-(4-bromanyl-2,6-dimethoxy-phenyl)-∼N-butyl-∼N-(cyclopropylmethyl)-2,7-dimethyl-pyrazolo[1,5-a][1,3,5]triazin-4-amine), but the binding site was predicted not as deep as in the experimental structure of this complex. Refinement of ligand binding sites prediction, in particular for the larger and more complicated targets, will be the key focus of further development of AstraBIND.

**Figure 1.**
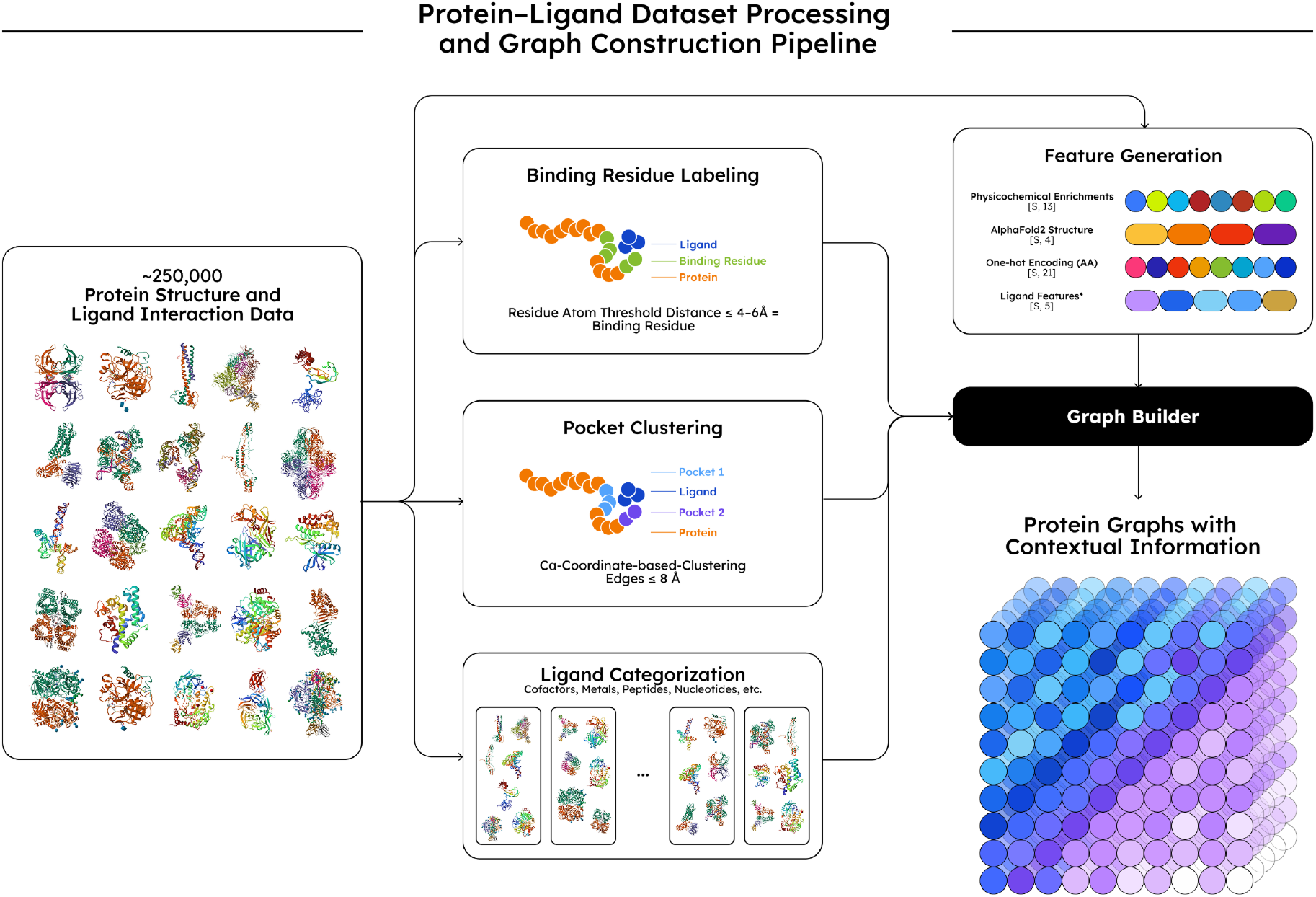
AstraBIND graph creation flow, showcasing how the interaction data is processed, structured, and enriched for before the training.

**Figure 2.**
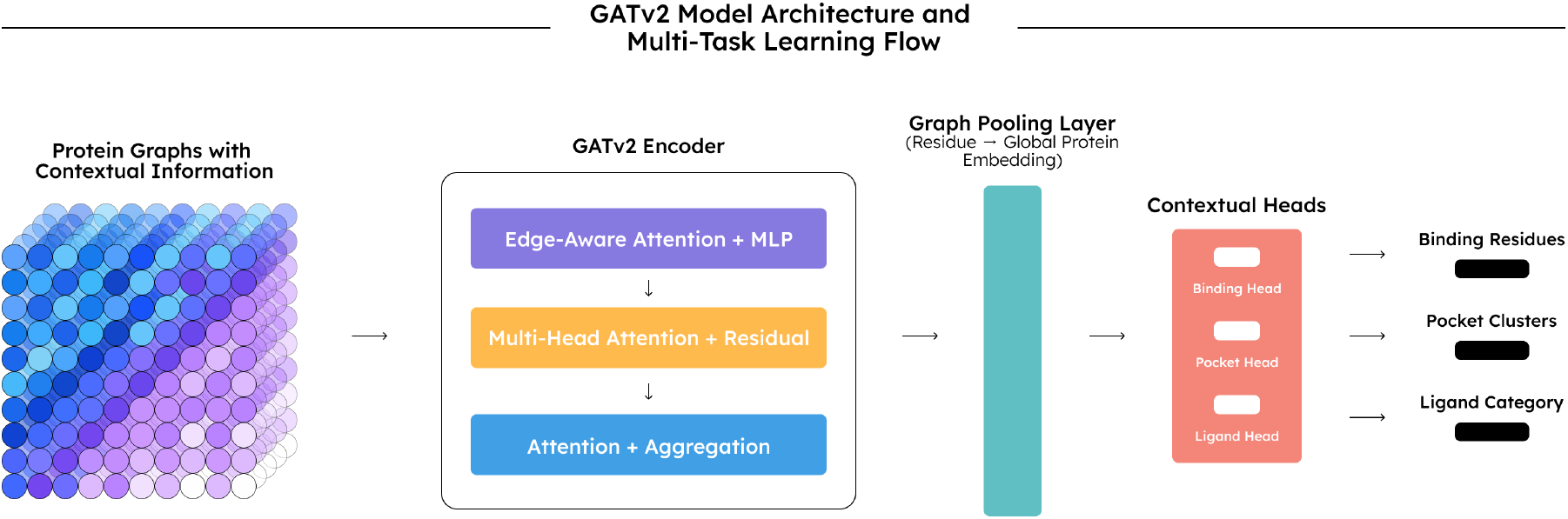
AstraBIND model’s training flow, processing the training data, and learning the binding site, pocket, and ligand patterns.

**Figure 3.**
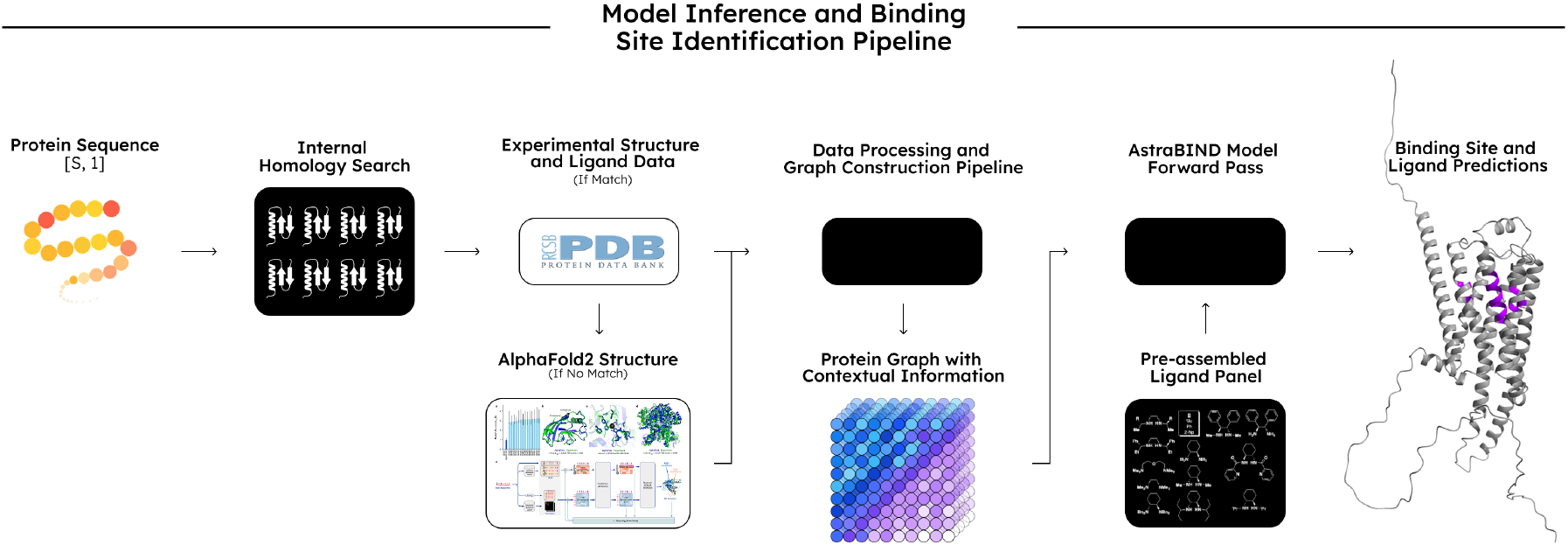
AstraBIND model’s inference flow, showing how the graph construction pipeline is integrated with model, and how the results are processed and represented on the user interface.

**Figure 4.**
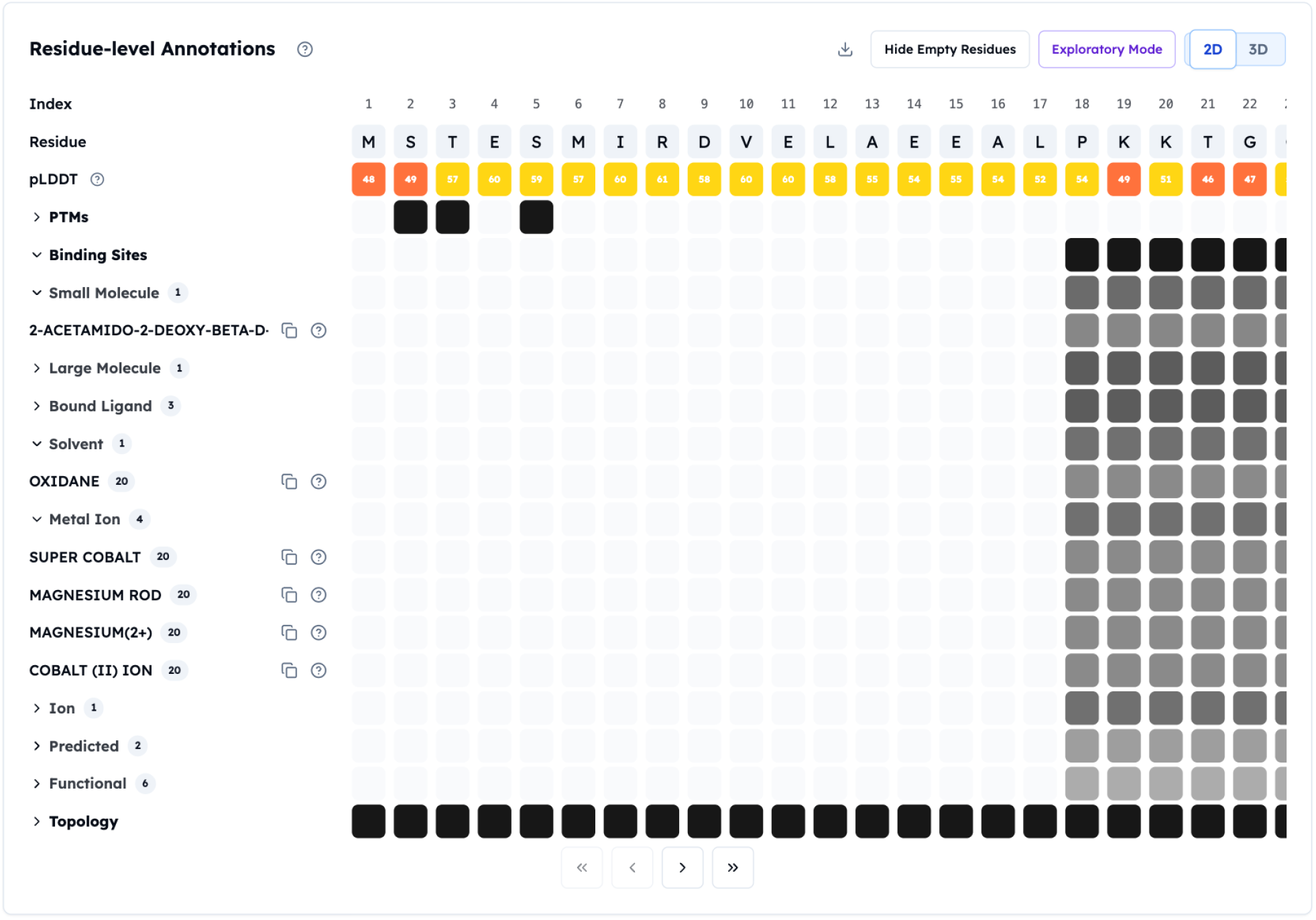
2D view of AstraBIND predictions for a representative target in the Orbion platform.

**Figure 5.**
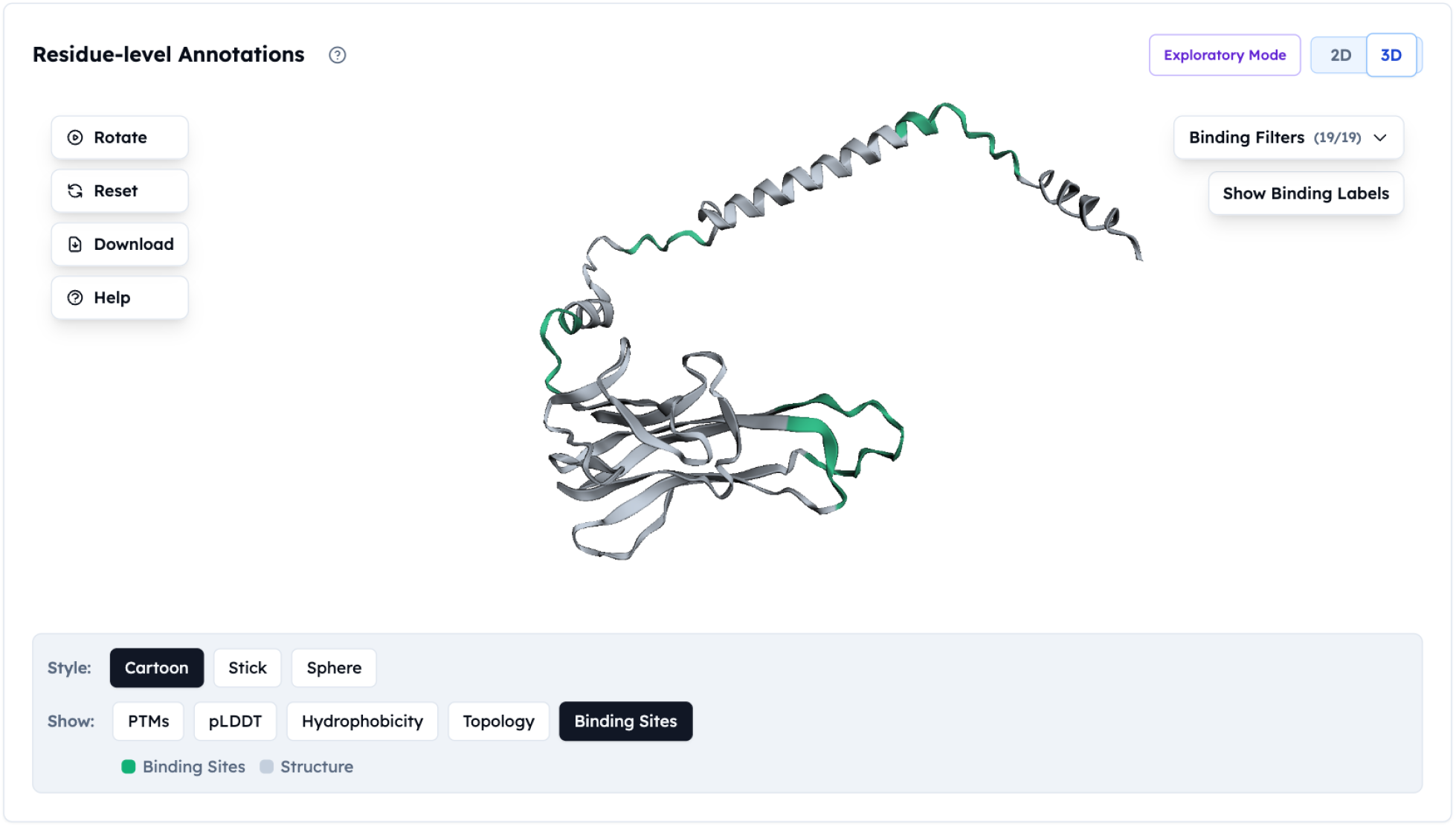
3D view of AstraBIND predictions using `3Dmol.js`. Predicted binding residues and pocket surfaces are overlaid on the structure.

**Figure 6.**
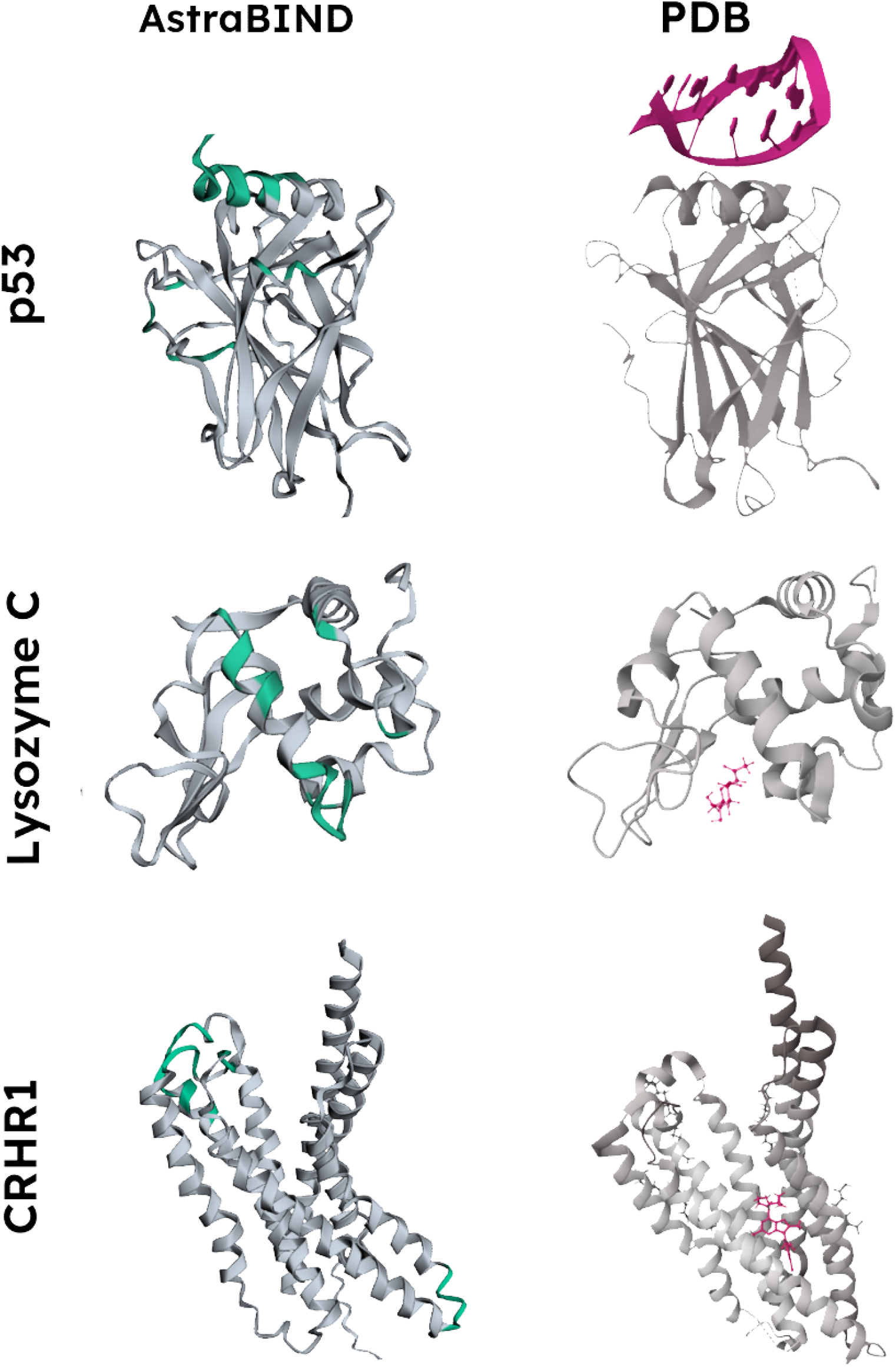
Comparison of AstraBIND binding residue predictions with experimental structures: p53 (UniProt P04637) predicted binding site vs X-ray diffraction structure of wild-type p53 (PDB 7B46); lysozyme C (UniProt P61626) predicted binding site vs X-ray diffraction structure with N-Acetyl-alpha-D-Glucosamine (PDB 7XF7); Corticotropin-releasing factor receptor 1 (CRFR1, UniProt P34998) predicted binding to 0JS ligand vs X-ray diffraction structure of complex with 0JS (PDB 8GTI). All proteins in this example are human. Predicted binding sites are shown in green and ligands are shown in pink. Other ligands and peripheral fragments are not shown.

## Discussion

A wide range of ML approaches have been proposed to tackle the problem of ligand binding site prediction.(14; 16) These can be classified into structure-based and sequence-based. Structure data almost always contributes to more accurate results compared with relying on sequence alone as input, however it also requires notably higher computational expenses.(14) In the past few years, several sequence-based models have approached structure-based methods in result accuracy thanks to the incorporation of protein language model (PLM) embeddings, which provide some of the amino acid context information without the need for explicit structure input.(14) Another consideration is that computer-generated structures (e.g. with AlphaFold) are becoming increasingly more reliable, meaning that structure information can be obtained even for novel proteins, or proteins without a known holo (ligand-bound) conformation structure.(16)

In the last few years, multiple models have been published using starkly different methods (Table 2). Even in the past two years, both models requiring structure information and structures relying on sequence input only have been made available. There is also a broad variety of geometry logic approaches: volumetric (voxel-based segmentation of a 3D-structure similar to a 3D-image), surface (segmentation into voxels or point clouds) and residue-level (e.g. distance graph) methods have all been used, each presenting their own advantages and drawbacks.

**Table 2.**
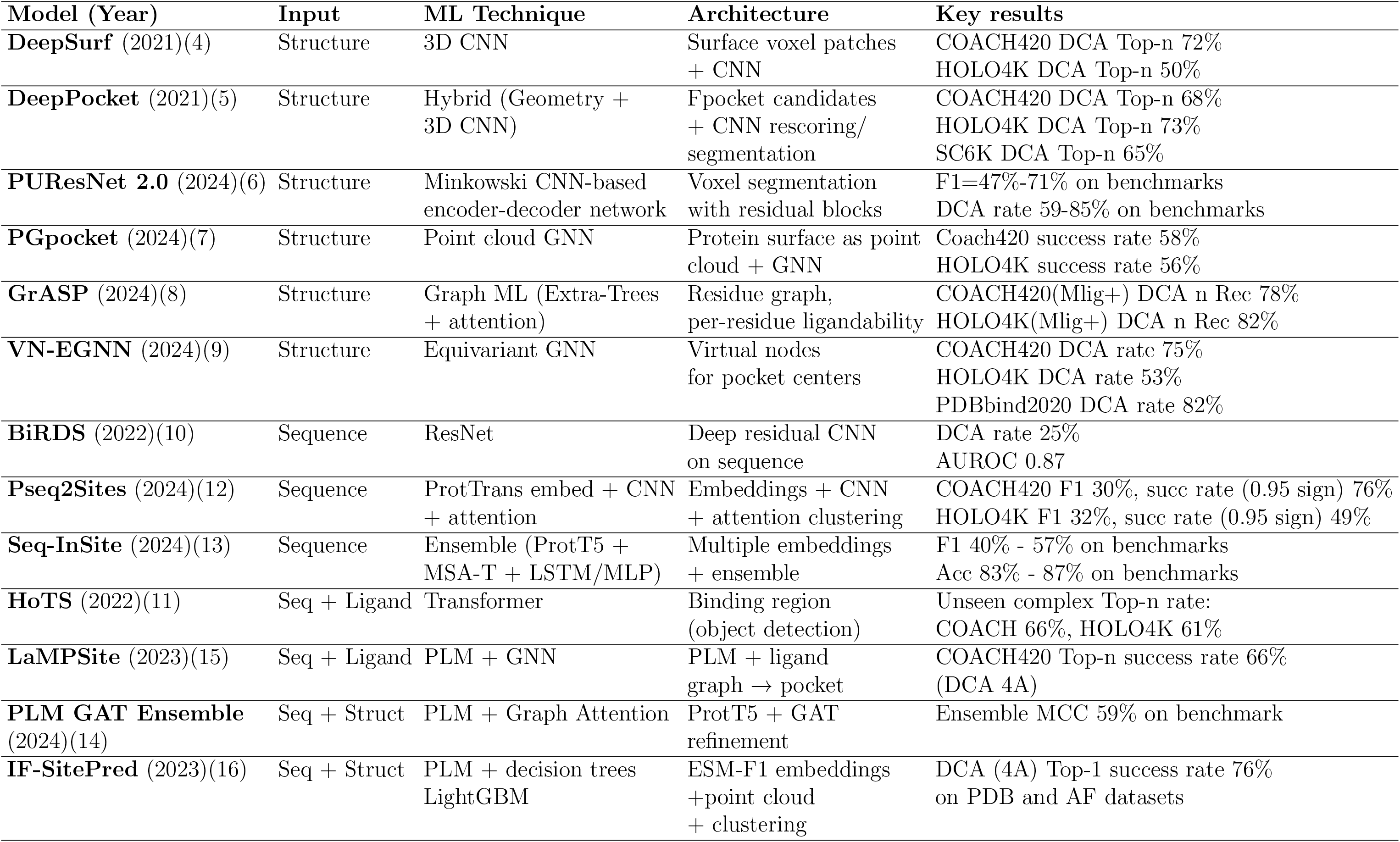
Key models for ligand binding prediction published in 2021 or later.

It is difficult to compare the performance of recent models directly, due to different approaches, benchmarks and metrics reported (see Table 2). However, as demonstrated by the lack of consensus on the best approach, all models had their drawbacks, and no single method demonstrated a substantial enough advantage in results over others.

Lack of consensus among recently published models indicates that ligand binding prediction tasks need to be considered in context, in particular with regard to the desirable trade-off between accuracy and speed. The Astra family of models is designed to provide actionable insights to scientists working with proteins in the lab, and need to process multiple protein and protocol iterations, in particular in mutation assessment tasks. For this reason, AstraBIND was constructed as a residue-based model (to provide sufficient sensitivity to small point mutation changes) but accounted for the 3D structure effectively via graphs and embeddings.

Combined with the high-quality training data, this approach produced a model that shows state-of-the-art performance at the high speed of just a few minutes per protein. It is also interesting to note that in some case studies, the model prioritised artificially strong ligands over natural ligands at high thresholds, which indicates some structure-based affinity evaluation, despite the fact that AstraROLE was not trained on affinity data. Although this cannot be used for numerical affinity predictions, it nonetheless serves as a sense check of the model predictions, and indicates that lower detection thresholds may be needed for popular drug targets that have artificial high-affinity ligands.

The model has several limitations. Although the AstraBIND model screens the protein against a large library of ligands, it is still possible that the protein’s ligand is not included; if the ligand scan returns null results, the model still functions using geometry alone, but this may result in lower prediction reliability. Structure dependency inference utilises experimental PDB data where possible, using AlphaFold2 predictions as a fall back; in the latter case, quality of structural predictions influences the AstraBIND output.

## Conclusions

AstraBIND results confirm that a spatially-aware graph infrastructure can support meaningful predictions for ligand binding sites, leveraging structural information to make effective evaluations. This is an essential step towards better protein understanding, underlying the generation of more advanced practical suggestions, such as laboratory protocols and stabilising mutations.

